# Investigation of a New Superhydrophobic, Oleophobic-Coated Biliary Duct Stent and its Preventive Effect on Biliary Mud in Vitro

**DOI:** 10.1101/708768

**Authors:** Wei Xing Zhang, Ya Li Ji, Yue Qin Qian, Xin Jian Wan, Zheng Jie Xu

**Author notes:** Correspondence author: Xin Jian Wan, MD, Department of Gastroenterology, Shanghai Jiao Tong University Affiliated Sixth People’s Hospital, Shanghai 200233, China. Co-correspondence author: Zheng Jie Xu, MD, Department of Gastroenterology, Xinhua Hospital Affiliated to Shanghai Jiao Tong University School of Medicine, Shanghai 200092, China. If you would like to chat with an author of this article, you may contact Dr Wan at or Dr Xu at. Each author’s contribution to the following criteria for authorship as follows: Wei Xing Zhang for main experimental operations, drafting of the article and critical revision of the article for important intellectual content, Ya Li Ji for coated stent and characteristic performance detection,Yue Qin Qian for bile collection, Xin Jian Wan for conception and design and final approval of the article,and Zheng Jie Xu for statistical analysis of data.

## Abstract

**Background and Aim:** A new type of superhydrophobic, oleophobic-coated biliary stent was created, and its characteristics and preventive effect on biliary mud deposition in vitro are studied herein.

**Methods:** The observational experiment included a bare stent group and coated stent group, with 10 stents per group. The groups were used in a model of the extracorporeal biliary perfusion system with bacterial infection, and the experiment was terminated when the stent was completely blocked. Changes of bile characteristics before and after the experiment, patency time, and amount of bile sludge deposition were compared between the groups. The t-test and analysis of variance were used to analyze the data.

**Results:** In the bare stent group, contact angles of bile were about 80.4 degrees and 79.8 degrees before and after the experiment, respectively. There was no change in the effect of bile aversion after the experiment between the groups (P < 0.05). In the coated stent group, contact angles of bile changed from 143.3 degrees before the experiment to 135.7 degrees after the experiment; the effect of the coating decreased significantly after the experiment (P > 0.05). The patency times were 8 weeks and 23 weeks in the bare and coated stent groups, respectively (P > 0.05). Depositions of biliary mud were 47 mg and 13 mg in the bare and coated stent groups, respectively (P > 0.05).

**Conclusion:** The superhydrophobic, oleophobic-coated biliary stent can repel bile better and play a role in preventing bile sludge deposition in the extracorporeal biliary perfusion system.

## 1 Introduction

Endoscopic plastic biliary stent implantation^[1,2]^ is the most commonly used method for treating benign, malignant biliary strictures. Because of the stent’s small expansion diameter, it is easy to cause restenosis due to biliary mud deposition within a short time. The stent often needs to be replaced repeatedly under endoscopy, which causes pain to patients, increases the cost of treatment, and increases the risk of complications^[3]^. Therefore, it is an urgent problem to prevent the deposition of biliary mud in stents and prolong the patency time of plastic stents as much as possible.

Water droplets that fall on lotus leaves will form a nearly spherical white, transparent water ball that rolls back and forth without infiltrating the lotus leaves, i.e., the “lotus leaf phenomenon.”^[4,5]^ Some scholars have been inspired to develop superhydrophobic, oleophobic coatings based on bionics principles. Since the coatings have hydrophobic and oleophobic properties, they can repel water and oil^[6,7]^.

Because bile^[8]^ contains a large amount of water as well as bile pigments, bile salts, cholesterol, lecithin, fatty acids, inorganic salts, and other ingredients, our research team designed a superhydrophobic, oleophobic coating, which is intended to be used on the inner and outer walls of a bile duct stent. It was assumed that since the coating has a bile-repellent effect on polar groups (e.g., water, ions, and polar amino acids) and non-polar groups (e.g., lipids and non-polar amino acids), it would repel bile. Bile mud in bile cannot be deposited on the inner wall of the bile duct stent; thus, a superhydrophobic, oleophobic coating might prevent the deposition of bile mud. This study aimed to investigate the characteristics and preventive effect of a superhydrophobic, oleophobic-coated biliary duct stent on biliary mud deposition in vitro.

The study was approved by the experimental centre of Shanghai General Hospital.

## 2 Methods

### 2.1 Materials

#### 2.1.1 Preparation of the coated biliary stent

Synthesis of fluorine-containing copolymers: Use 3-(trimethoxymethylsilyl) propyl methacrylate, perfluoroalkyl ethyl acrylate, and dimethylaminoethyl methacrylate as reactants, add those reactants in a ratio of 1:8:16 (according to the molar ratio) in a 100-mL flask, add methyl isobutyl ketone as reaction solvent, and add an appropriate amount of an initiator, azodiisobutyronitrile, (0.5% mass of the total substance) into the flask. Remove nitrogen for 30 minutes and then heat it to 65 degrees Celsius for 18 hours. After the reaction is completed, the above solution is precipitated with a large amount of n-hexane, and the precipitate is a fluorinated copolymer. Then the copolymer is cleaned several times with n-hexane, cooled and dried, and stored in a dryer.

Preparation of coated biliary stent: A plastic bare stent (7.5-French, 5 cm) was placed in acetone, ethanol, and distilled water for 15 minutes; it was dried in a vacuum drying box at 60 degrees Celsius for 24 hours; and then it was disinfected with ethylene oxide. Next, a certain amount of a fluorine-containing random copolymer was evenly dispersed in anhydrous toluene to form a mixed solution. The bare stent was clamped by sterilizing a medical clip. The clean plastic biliary stent was immersed in the aforementioned mixed solution for 30 minutes and then it was removed and dried at 60 degrees Celsius for 4 hours. Finally, the unreacted fluorinated copolymers were washed with anhydrous ethanol and distilled water and dried at 60 degrees Celsius. The dried coated biliary stent was sterilized with ethylene oxide and packaged for reserve. Ten coated stents were created (Figure 1).

**Fig. 1.**
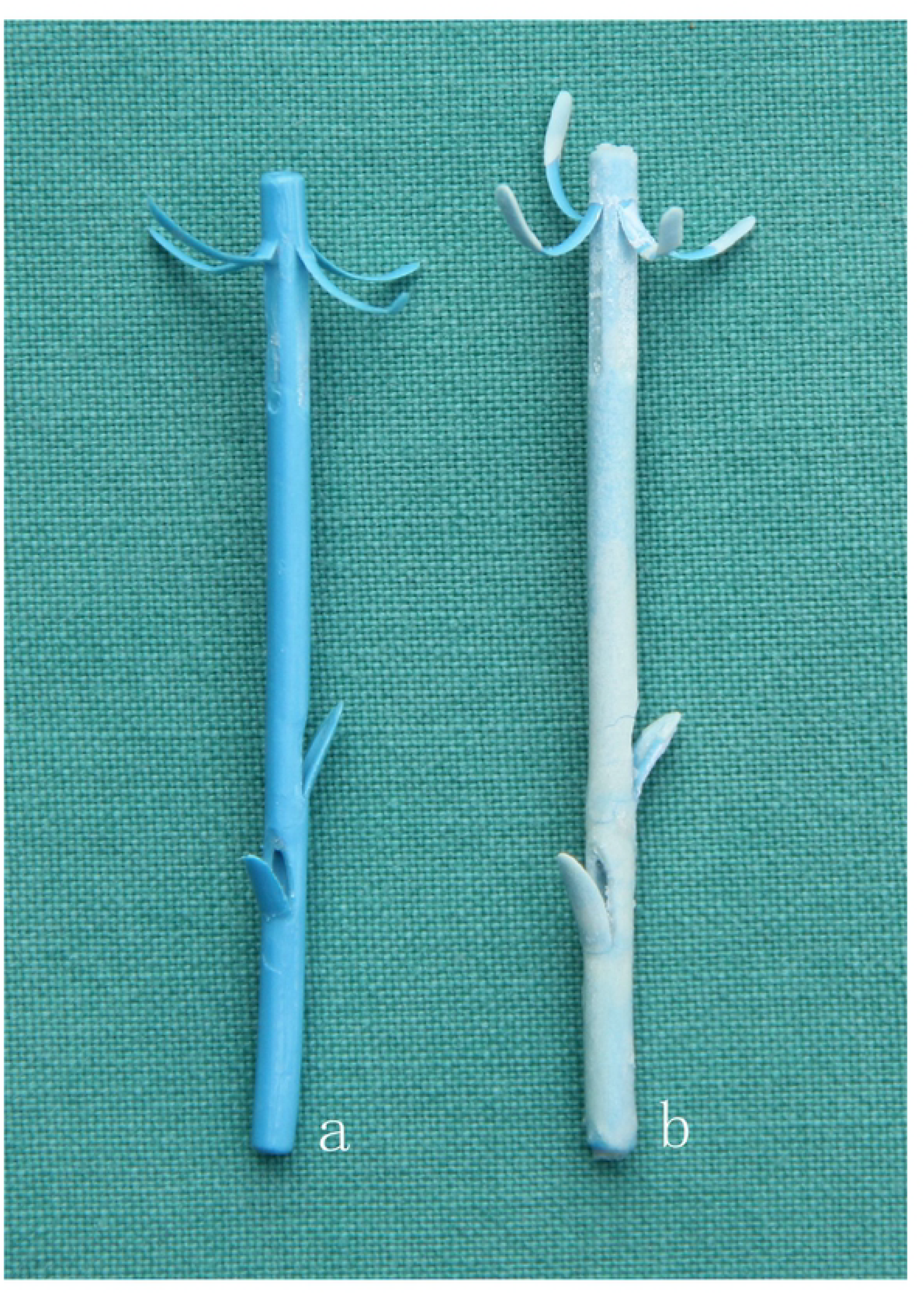
Biliary stent (a: bare stent, b: superhydrophobic, oleophobic-coated stent)

Assessment of the coated biliary stents: All stents were measured by two persons separately. If the data measured by two persons were inconsistent, we checked whether the data were documented incorrectly. If there were no documentation errors, the thickness of the stents was re-measured. The stent coating was closely attached to the plastic stent, which was uniform in thickness and not easy to peel off; otherwise, it was discarded.

Inspection: The coating thickness was measured at three points: 1 cm away from both ends and in the middle by micrometers. The average value was recorded. If the average was 0.1 ± 0.01 mm, the stent was re-coated or discarded.

### 2.2 Experiment

#### 2.2.1 Characterization and performance testing

Observe the coating with naked eyes: The shape of bile on the bare stents and coated stents was observed. We determined whether the bile formed spherical bile beads or rolled freely without infiltrating the bare stents and coated stents.

Scanning electron microscopy test: Scanning electron microscopy (SEM) (SU8010, Hitachi, Tokyo, Japan) was used to observe the whole surface of the bare stent and coated stent.

Contact angle measurement: The JC2000C1 contact angle measuring instrument (Shanghai Zhongchen Digital Equipment Co., Ltd., Shanghai, China) was used to measure the contact angle of bile in the inner wall of the bare stent and coated stent. The change of the contact angle over time was observed and measured five times, and the average value was recorded.

#### 2.2.2 Perfusion test using a model of the biliary perfusion system

The experimentation area of the bile duct perfusion system was numbered, and 10 bare stents and 10 superhydrophobic, oleophobic-coated stents were placed. After disinfection with an ultraviolet lamp, the porcine bile was added into the liquid box. *Escherichia coli* was added to the pig bile. The bacterial content was 1-5 × 10^6^. After 2 hours of culturing at 37 degrees Celsius, the bile was passed through the valve via the shunt and then into the sample. The valve was regulated so that the amount of bile collected per stent was about 1000 ml per day, which is similar to the amount of bile discharged by the liver daily (Figure 2).

**Fig. 2.**
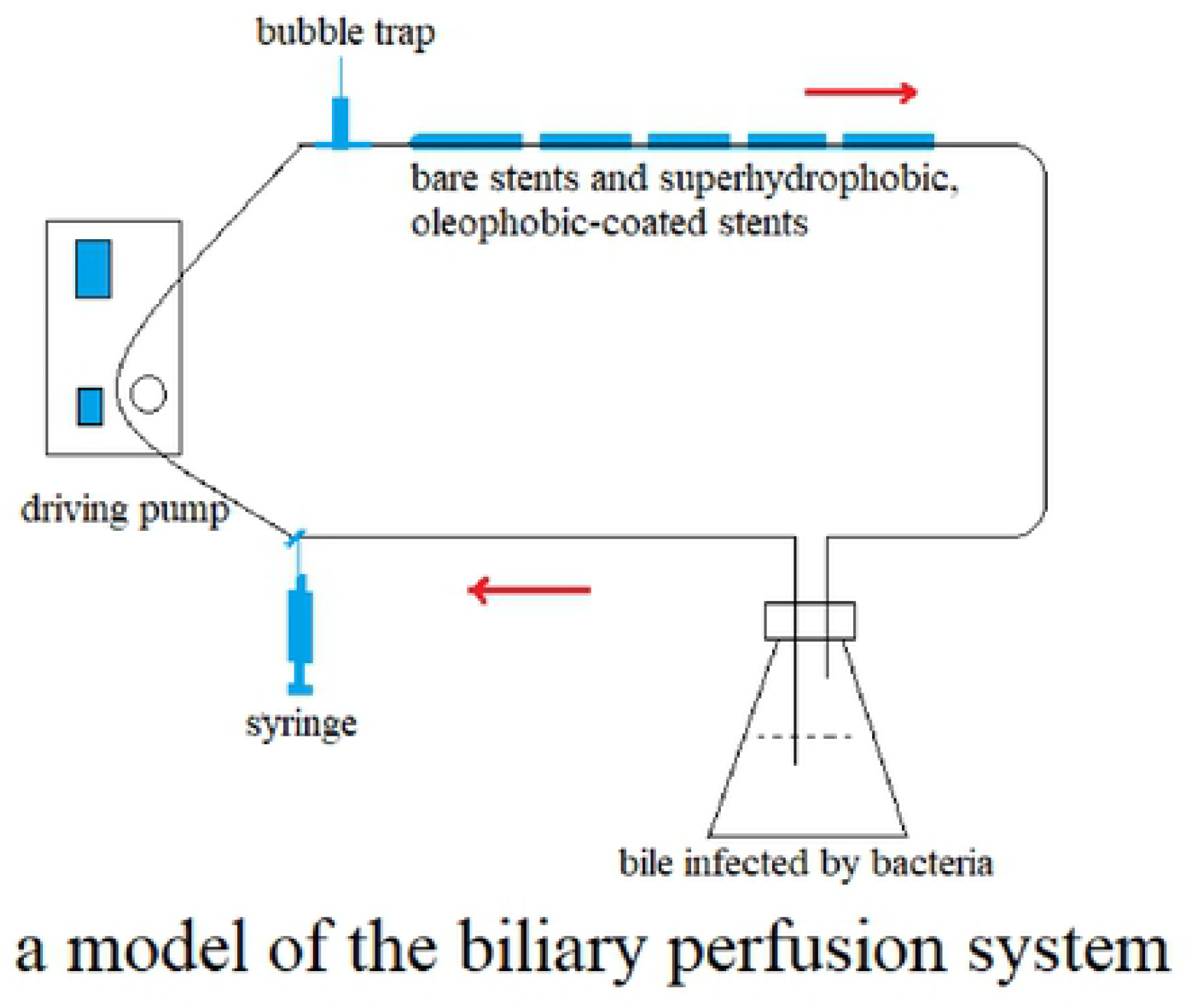
Experiments with the stents in the in-vitro biliary perfusion system model with bacterial infection

#### 2.2.3 Observation items

After the stent was completely blocked, the inner and lateral walls of the bare stent and coated stent were observed under naked eyes and electron microscopy. The contact angle, distribution of biliary mud, and weight of amount of biliary mud were measured.

Observation of the anatomical structure of the inner wall of the biliary stent: The bile was washed gently under normal saline, and changes of the wall of the stent were observed after it was air dried at room temperature. The bile duct stents were dissected cross-sectionally to observe the distribution of bile mud. The whole surface structure of the bare stent and coated stent were assessed to determine whether it was smooth or rough, whether the structure was damaged, whether the coating was peeled or damaged, and whether the bile formed spherical bile beads or rolled freely without infiltrating the stent.

Scanning electron microscopy test: SEM was used again to observe the whole surface of the bare stent and coated biliary stent (Figure 3).

**Fig. 3.**
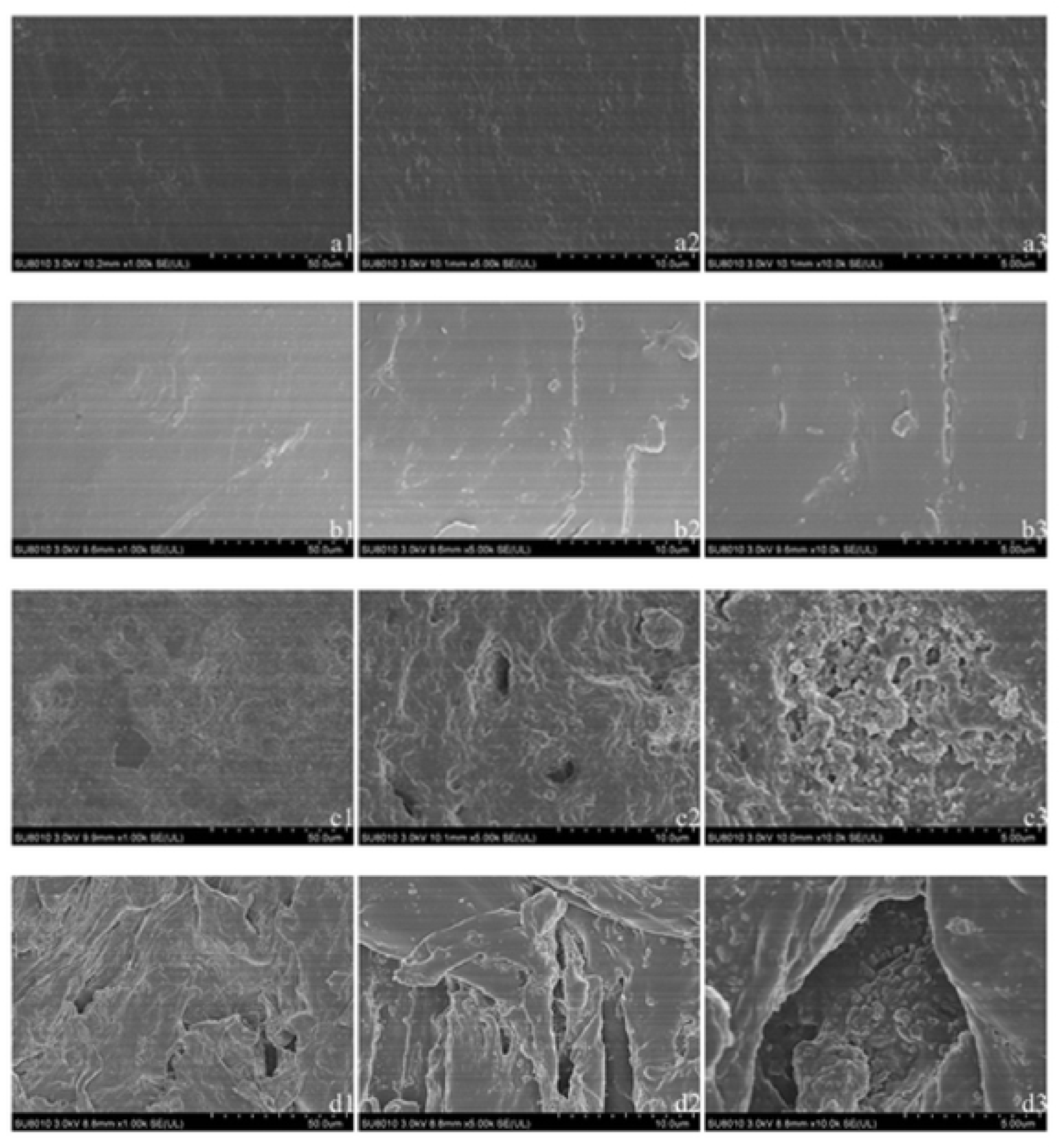
Electron microscopic photographs (bare stent: a1 ×2000 times, a2 ×10 000 times, a3 ×20 000 times before the experiment; b1 ×2000 times, b2 ×10 000 times, b3 ×20 000 times after the experiment and coated stent: c1 ×2000 times, c2 ×10 000 times, c3 ×20 000 times before the experiment; d1 ×2000 times, d2 ×10 000 times, d3 ×20 000 times after the experiment)

Measurement of the contact angle: The contact angles of bile for the bare stent and coated stent were measured. Changes of the contact angles over time were observed. The average values were measured five times (the minimum measurement value was 0.1 degrees) (Figure 4).

**Fig. 4.**
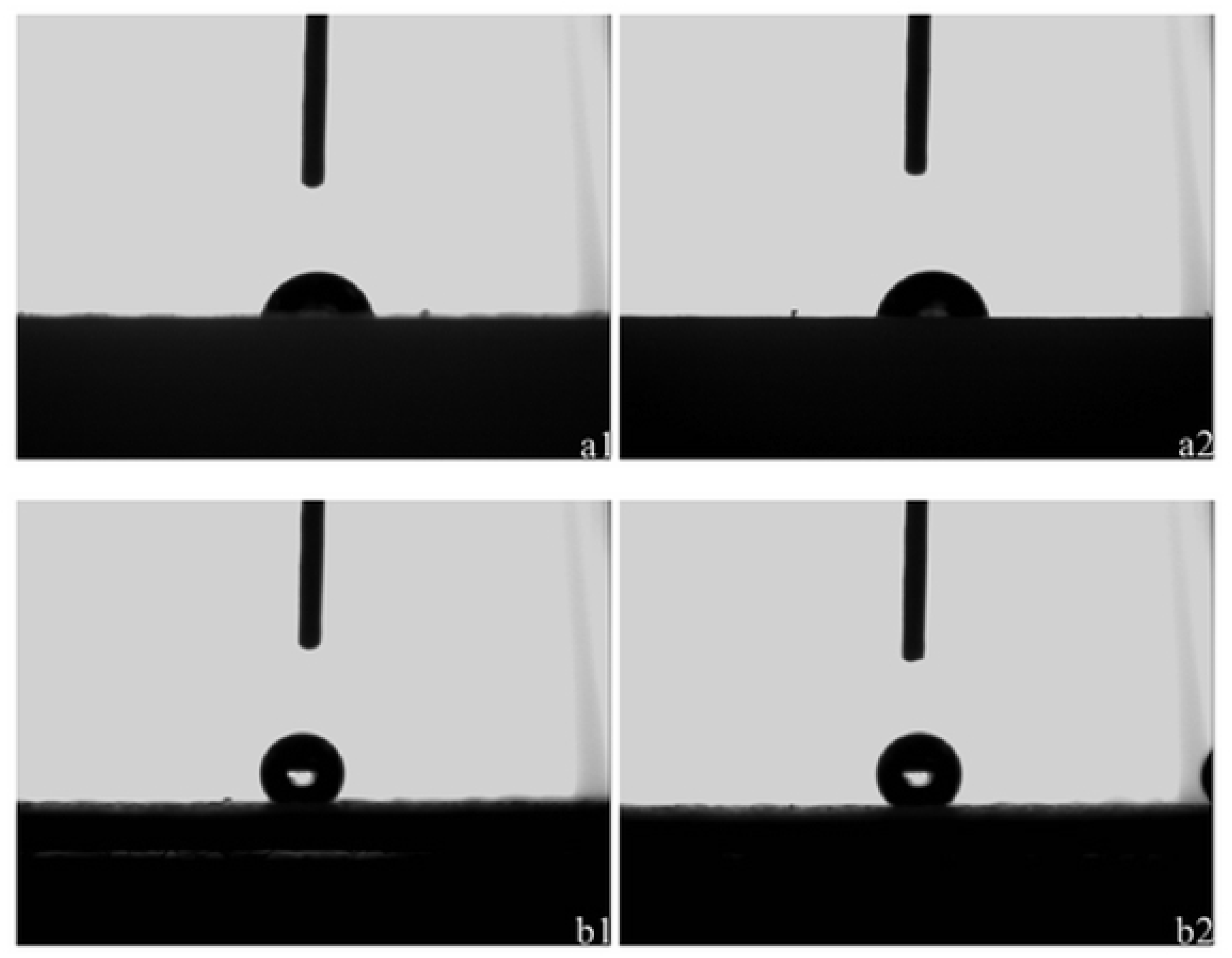
Bile contact angle measurement (bare stent: a1 before the experiment, a2 after the experiment; coated stent: b1 before the experiment, b2 after the experiment)

Quality of bile mud: After all the bile mud was removed, it was weighed using an analytic balance three times, and the average value was recorded. Then the amount of bile mud was calculated by the equation(the quality=sum of three measurements/3)(the minimum measurement value was 0.1 mg).

#### 2.2.4 Follow-up time and assessment of outcome

Time of stent patency: The stent was removed every weekend to observe its shape and whether it was obstructed. To simulate the human bile duct pressure, hydrostatic pressure of 50 cm was maintained, and bile was dripped slowly from one end of the stent to the other. If the bile could flow to the other side, the stent was considered to be not completely blocked, and the perfusion test was continued. If the bile could not flow to the other side, the stent was considered to be completely blocked, and the stent patency test was terminated. The time at which the stent was completely blocked was the patency time. The follow-up time until all stents were obstructed was calculated in weeks.

#### 2.2.5 Statistical analysis

The t-test and analysis of variance were used to compare data between the groups. The results are expressed as 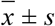. SPSS version 17.0 statistical software (IBM Corp., Armonk, NY, USA) was used to analyze the data. P < 0.05 was statistically significant.

## 3 Results

### 3.1 Characteristics of bile on the coated stent before and after the experiment

Naked eye observation: Before the experiment, the color of the bare stent was blue, its surface was smooth, and bile beads, which could not roll freely, formed in a semicircle on the bare stent. After the experiment, the color of the bare stent became black, and its inner wall became filled with bile mud. After removing the bile mud, some proteins and mucus adhered to the inner wall of the stent, its surface was still smooth, and bile beads, which could not roll freely, formed in a semicircle on the bare stent (Figure 4(a1, a2)).

Before the experiment, the color of the superhydrophobic, oleophobic-coated stent was grey, rough, and wrinkled, but the structure was intact and surface was relatively flat. Bile formed spherical bile beads on the coating, and it could roll freely without infiltrating the coating. After the experiment, the color of the coated stent was grey-black, its upper wall was filled with bile mud, and middle and lower end of the coating deteriorated. Part of the superhydrophobic, oleophobic coating fell off and broke, especially on the upper end. The surface of the coating that remained was relatively flat. Bile formed spherical bile beads on the intact coating, and it could roll freely and not infiltrate the coating.

SEM test and analysis: Before the experiment, the surface structure of the bare stent was neat and flat without protrusion, depression, or other surface features (Figure 3(a1, a2, a3)). After the experiment, the surface structure of the bare stent was still neat and flat without obvious protrusion, depression, or other surface features, and there was no significant difference from before the experiment (Figure 3(b1, b2, b3)).

The surface morphology of the coating was observed by SEM. Many tiny papillae were present on the surface of the coating. The average size of the papillae was about 5-10 microns, average height was approximately 10-15 microns, and average spacing was about 20 microns. Among these small papillae, there were some larger papillae, with an average size of about 50 microns. They also comprised 5-15 micron-sized micro-processes. The top of the mastoid process was flat and slightly depressed in the center. This mastoid structure is difficult to detect with the naked eye and ordinary microscopy, and it is often referred to as a multi-nanometer and micro-scale ultrastructure. The large and small papillae and protrusions on the lotus leaf-life surface were like hills. The depressions between the hills were filled with air. Thus, a very thin layer of air was formed on the lotus leaf-like surface, which was only a nanometer thick. The ability to repel bile was strong (Figure 3(c1, c2, c3)).

After the perfusion test, the micro-morphology of the coating was observed by SEM. The small papillae and large papillae on the upper surface of the coating had fewer mutations, and the average size, height, and spacing of the papillae became larger. In some places, the papillae were damaged or disappeared. The top of the mastoid process was flat with central depression, and the ultrastructure was destroyed. The depression was filled with less air, and the ability to repel bile was weakened. The surface of the coating became very rough, and the wrinkles worsened. The difference before and after the experiment was obvious (Figure 3(d1, d2, d3)).

Contact angle test: The contact angle between the bile and bare stent were about 80.4 degrees before the experiment and 79.8 degrees after the experiment. There was no significant difference in the effect of bile aversion after the experiment (P < 0.05); that is, over time, the contact angle did not change significantly, and the surface of bare support was stable (Figure 4(a1, a2)).

The contact angles of bile coated with superhydrophobic, oleophobic coating were about 143.3 degrees before and 135.7 degrees after the experiment. The effect of bile coated with superhydrophobic, oleophobic coating was significantly weakened after the experiment (P > 0.05). However, the bile had a good spherical structure on the coating, and the coating still exhibited good, stable superhydrophobic and oleophobic properties (Figure 4(b1, b2)).

### 3.2 Relationship between the biliary stent and stent patency time

In the biliary perfusion system, the patency times of the bare stent and coated stent were about 8 weeks and 23 weeks, respectively. Thus, the patency time of the coated stent was about 15 weeks longer than that of the bare stent.

### 3.3 Amount of biliary mud in the stents

The distribution of biliary mud in the bare stent was uniform, and that in the upper end, middle, and lower end of the stent was uniform; additionally, obstruction occurred uniformly in the stent lumen. The internal cavity of the coated stent was not uniform. The biliary sludge was mainly distributed in the upper end of the stent, with less in the middle and lower end. The obstruction mainly occurred in the upper end of the coated stent. These differences were also reflected in the degree of damage and shedding of coatings, which is consistent with the damage and shedding of coatings.

The total amount of bile sludge deposited in the bare stent group when the stent obstructed was about 47 mg and that in the coated stent group was about 13 mg. There was a significant difference between the groups (P > 0.05).

## 4 Discussion

The β-glucuronidase produced by a bacterial infection of biliary tract can decompose bilirubin-glucuronide diester, produce bilirubin, and bind with ionic calcium, which leads to the deposition of calcium bilirubin, and then to the formation of biliary mud and bile duct stones^[9,10,11]^. A small amount of fibronectin, collagen, fibrin, and immunoglobulin A in bile accelerates the biliary mud covering the stent surface. Bile mud deposits gradually in the stent, which eventually leads to stent occlusion. In our investigation of a fabricated superhydrophobic, oleophobic-coated biliary stent, we found that the fluorinated copolymer can form a coating that is firmly bound to the base, and the coating has good superhydrophobic and super-thin oil properties.

In addition, the contact angle of the superhydrophobic oil-thinning coating to bile showed a good bile-repellent effect. A barrier film was formed between the bile and coating. The water, ions, and lipids in the bile did not adhere to the coating. Because of the surface tension of bile, the shape of the bile on this rough surface was nearly spherical, and the bile could roll freely on the surface. Bile could not adhere to the coating, and mucin and mud in the bile were separated by the coating. It was not easy for bile to deposit on the coating and form mud. At the same time, because the bile contains a large amount of water and bile pigments, bile salts, cholesterol, lecithin, fatty acids, inorganic salts, and other components, the coating had a bile-repellent effect. It also isolated polar groups and non-polar groups in the bile, and the bile sludge in the bile could not be deposited in the bile duct branches. The inner wall of the stent prevented the deposition of bile mud.

We coated the inner and outer walls of the bile duct stent with a superhydrophobic, oleophobic coating, which made the inner and outer walls of the duct that contacted the bile have a protective film that was formed by a fluorine-containing nano-polymer. The film had good superhydrophobic and oil-draining properties, isolated the inner and outer walls of the bile and bile duct stent, prevented the deposition of bile mud on the stent, and prolonged the time of catheter replacement.

Generally, the stronger the bond between the coating and stent, the better the mechanical properties of the coating itself, the better the bile-repellent effect of the coating, the longer the time of stenting, and the stronger the effect of preventing bile mud deposition. Although the bare stent itself has certain inertia and stable performance, the superhydrophobic oil-thinning coating on the bare stent will break and fall off under the pressure of bile, and the surface of the coating will sustain wear and tear. The surface structure of some of the coatings changed, some protrusions disappeared, depressions widened, contact angle decreased, and bile-repellent performance weakened due to wear and tear. Further, part of the coating cracked and the leaky bracket of the bare stent lacked a bile-repellent effect due to wear and tear, which resulted in the deposition of bile mud.

Therefore, some of the superhydrophobic, oleophobic coating on the biliary stent peeled off during the experiment. Additionally, the coating became damaged under the pressure of bile, which weakened the bile-repellent effect of the coating over time, causing deposits of the bile duct at a later stage. The contact angle of the bare stent did not change much, but the contact angle of the coated stent decreased and subsequent deposits of bile mud occurred. After coating, the inner wall of the coated biliary stent becomes smaller because of the thickness of the inner wall coating. This factor is also the reason why the coated biliary stent was easy to obstruct, and the amount of deposited biliary mud was less with the bare stent than with the coated stent. Because of the presence of some mucin in bile, which are sticky and do not flow easily, bile duct obstruction occurs after the deposition of bile mud. All the obstructions we observed occurred mainly at the upper end of the coated stent, and the deposition of bile mud in the bare stent was more uniform than that in the bare stent.

If physicians need to prolong the patency time of a biliary stent, the coating technology needs to be improved in order to make it more firmly adhere to the stent surface and not easily peel off. Alternatively, the stent surface itself needs to be superhydrophobic and oleophobic, or the inner diameter of the stent needs to be expanded.

At present, results of the coated stent were observed in an extracorporeal biliary perfusion system. It needs to be investigated whether the coated stent can achieve the same effect in vivo. Because bacterial infection in vivo is generally not obvious, the body secretes fibronectin, collagen, fibrin immunoglobulin A, and other inflammatory mediators. These substances may have some effect on the deposition of bile mud and the biological safety of this new coated stent in vivo. This is the focus of our next research study.

This study has potential limitations. The effect estimates in the model are based on observational studies. They are therefore subject to biases and confounding that may have influenced our model estimates. Firstly, the contact angle of coated stent is measured and sampled differently,because of the different uniformity of materials and the exfoliation degree of different parts. Secondly, the coating thickness of stent inwall and the size of stent cavity can not be measured, and the sample size is only 10, so there might be bias.Thirdly, under the natural air-drying condition, the quality of bile mud varys with drying degree and moisture content. so there might be bias in outcomes. To increase the sample size and improve the coating technology might reduce those potential bias.

### 4.1 Conclusions

The superhydrophobic, oleophobic-coated biliary stent can prevent the deposition of bile sludge in vitro. The longest patency time of the stent was about 15 weeks, and the total amount of bile sludge deposited in the bare stent group when the stent obstructed was about 47 mg and that in the coated stent group was about 13 mg. There was a significant difference between the groups.

## Conflicts of Interest

None

## Notes

DISCLOSURE: All authors disclosed no financial relationships relevant to this publication.

